# The *Drosophila* EGFR ligand mSpitz is delivered to cytoplasmic capes at sites of non-canonical RNA export on the nuclear envelope *via* the endosomal system

**DOI:** 10.1101/2020.07.14.203000

**Authors:** Floyd J. Mattie, Praveen Kumar, Mark D. Travor, Kristen C. Browder, Claire M. Thomas

**Affiliations:** Departments of Biology and of Biochemistry and Molecular Biology, The Pennsylvania State University, University Park, Pennsylvania, United States of America; Department of Nutritional Sciences, The Pennsylvania State University, University Park, Pennsylvania, United States of America; Department of Zoology, Government College for Women, Trivandrum, India; Genentech Inc., 1 DNA Way, South San Francisco, California 94080, USA

**Keywords:** Cell Nucleus, Nuclear Envelope, Protein Transport, Endosomes, Ribonucleoproteins, EGF Family of Proteins, Spectrin, *Drosophila*

## Abstract

Nuclear-cytoplasmic communication is not limited to nuclear pores, with both proteins and RNA using alternative routes between these compartments. We previously characterized cytoplasmic capes (large invaginations of the nuclear envelope in *Drosophila*), which are enriched for the membrane-bound EGF receptor ligand mSpitz, endosome-related organelles and ubiquitylated proteins. Closely associated with capes are groups of perinuclear vesicles resembling those seen at sites of RNP export *via* a budding mechanism. Here, we demonstrate that mSpitz delivery to capes requires passage through the endosomal system. We also show that capes are closely associated with sites of non-canonical RNP export as well as the dFrizzled2 receptor C terminal fragment, a core component of this export pathway. Video microscopy of glands in intact larvae indicates that cytoplasmic capes are stable structures that persist for at least 90 minutes without conspicuous growth. We further show that capes appear with the growth of the salivary gland rather than its developmental stage. Finally, we show that the large F-actin binding protein β_H_-spectrin, which modulates endosomal trafficking, as well as its partner α-spectrin are required for cape formation. Cytoplasmic capes therefore represent a subspecialization of the nuclear envelope where endosomal trafficking and RNP export are closely associated.

**Synopsis:** We further characterize large invaginations of the nuclear envelope called cytoplasmic capes in *Drosophila*. The EGF receptor ligand mSpitz is concentrated in capes and we show that it traffics to this compartment *via* endosomes. The presence of RNP and the dFrizzled2 receptor C-terminal fragment also indicates that non-canonical RNA export is concentrated at capes. *In vivo* imaging shows that capes persist for at least 90 minutes. Finally, the large F-actin crosslinker α/β_◻_-spectrin is shown to be required for cape formation.

**Abstract Figure:** 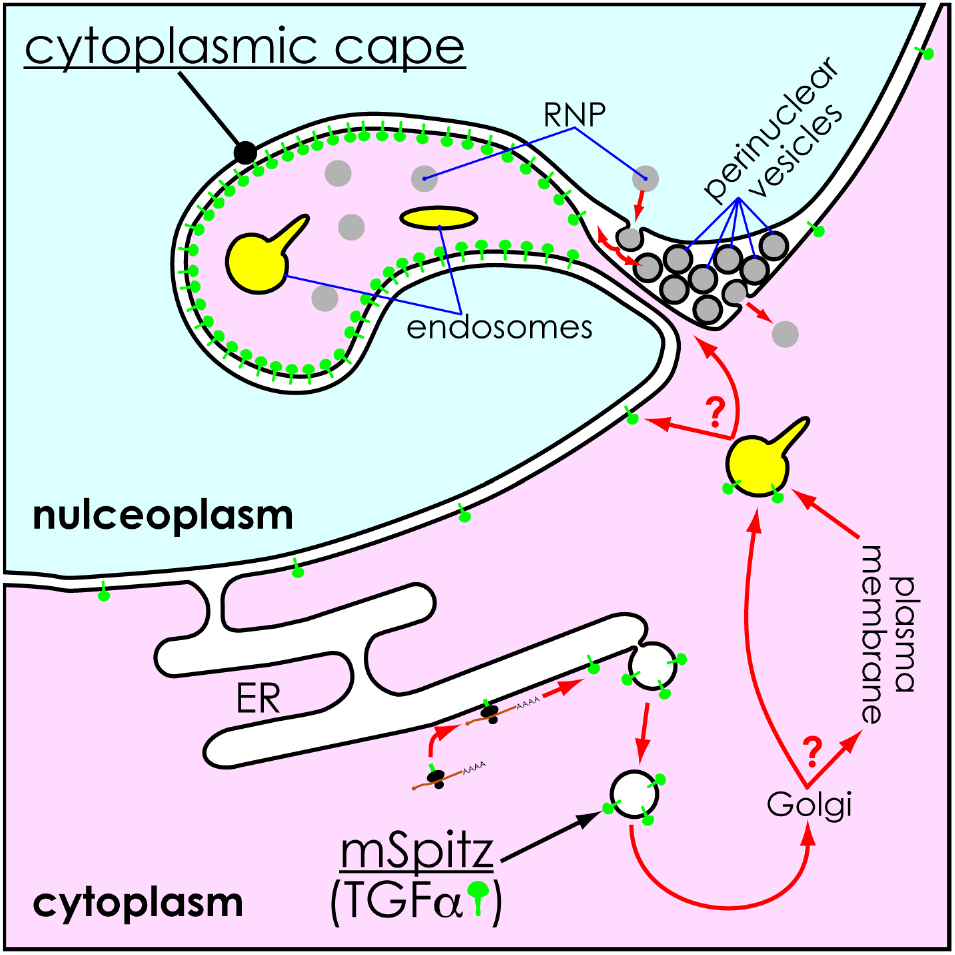

## Introduction

The routes by which the import and export of materials to the nucleus occurs are varied. Passage to and from the nucleoplasm through nuclear pores is increasingly well-understood ^1,2^. It is also increasingly evident that there are multiple routes by which proteins are delivered to the nucleus ^3^. A well-established route to the inner nuclear membrane (INM) is *via* lateral diffusion from the contiguous endoplasmic reticulum (ER) to the outer nuclear membrane (ONM), followed by passage to the INM at sites of contact in nuclear pores ^4^. In some cases proteins may first travel out to the Golgi before retrograde return to take advantage of this pathway. Joining this route are some proteins that are internalized at the plasma membrane and pass through the endosomal system ^5–7^. Recently, it has also been shown that some proteins endocytosed from the plasma membrane to the endosomal system are delivered to the nucleoplasm *via* fusion of nuclear-envelope associated endosomes directly with the ONM and passage through the Sec61 translocon in the INM ^8,9^.

Some membrane bound receptors end up in the nucleoplasm itself (probably in association with chaperones to mask their hydrophobic transmembrane domains), and here the Sec61 translocon appears to have an important role. Sec61 is best known for it’s role in extracting proteins from the ER for the ERAD protein surveillance pathway ^10,11^. However, it also plays a role in extracting membrane proteins from the ER for import to the nucleoplasm using importins to bring the protein through the nuclear pore complex ^12^ and on the INM where it directly extracts proteins into the nucleoplasm ^13–15^

RNA export mechanisms also exhibit diversity. Whereas RNA export through nuclear pores is well established, there is also a non-canonical RNA export (NCRE) pathway for large RNP’s that bypasses nuclear pores *via* budding from the INM followed by fusion to the ONM ^16^. RNP budding in *Drosophila*, from the INM is regulated by Wingless signaling *via* the dFrizzled2 receptor C-terminus (DFz2-C) that is cleaved and enters the nucleus to stimulate aPKC phosphorylation of Lamin C ^16^. Export of perinuclear vesicles from the INM also requires the activity of the AAA+ ATPase Torsin ^17^ and ESCRT-III^18^. Defects in this pathway result in trapping of specific RNA’s in the nucleus and prevent accumulation of RNP’s in the region of the neuromuscular junction. In humans, mutations in lamins may also affect this pathway and contribute to deficits associated with some laminopathies and/or dystonia ^16,17^.

While investigating the role of spectrin in endosomal transport in *Drosophila* we discovered that mSpitz, the membrane bound precursor of the EGF Receptor ligand sSpitz, strongly accumulates in nuclear envelope invaginations ^19^. These structures, called cytoplasmic capes 20, were originally seen in ultrastructural studies of *Drosophila* salivary gland nuclei^21^ and are distinguished from the nuclear reticulum in scale and morphology ^22,23^. We previously demonstrated that these distinctive structures are strongly associate with endosome-like organelles and contain ubiquitylated proteins ^19^. Associated with these structures were clusters of perinuclear vesicles ^19^, strongly resembling those containing mega-RNP’s in muscle nuclei ^16^, and probably related to RNA-containing ‘oval bodies’ in older references ^20,24,25^. In our initial investigations it is clear that the protein content of the capes is selective ^19^; however, just how proteins accumulate in capes has not been investigated, nor has their relationship to NCRE.

In this paper, we have investigated the route by which mSpitz is delivered to cytoplasmic capes and explicitly demonstrate a relationship between capes and NCRE. We show that the EGFR ligand precursor mSpitz accumulates in cytoplasmic capes after passage through the endosomal system. Several factors important for endosomal trafficking are important for the accumulation of mSpitz at this location, but none had a conspicuous effect on cape formation *per se*. This suggests that this protein accumulates in cytoplasmic capes either through retrograde transport *via* the ER or by direct transfer from nearby nuclear-envelope associated endosomal compartments. In addition, we use an improved *in situ* hybridization technique to show that most, if not all, cytoplasmic capes have associated sites of NCRE and that these structures are also positive for DFz2-C, a core component of these particles. We go on to show that, in contrast to previous studies, capes are not developmentally regulated strictly by developmental stage (instar), but appear as the salivary gland reaches a critical size threshold. We also examined nuclear cape dynamics in whole larvae and find that they are persistent structures lasting at least 90 minutes, limiting some models for their formation. Finally, we find that β_H_-spectrin, a large F-actin binding protein and modulator of the endosomal system, is required for efficient cape formation along with its molecular partner α-spectrin.

## Results and Discussion

We previously found a strong association between cytoplasmic capes and the presence of compartments containing endosomal markers and ubiquitylated proteins ^19^. These results suggested a possible connection between this sub-compartment of the nuclear envelope, cargoes such as mSpitz∷GFP, and endosomal transport. To further investigate this relationship we utilized inducible RNA hairpin constructs to knockdown various key components of the endomembrane system along with immunofluorescent staining for Lamin C to visualize capes and expression of mSpitz∷GFP as a representative cargo that accumulates in these structures. Out of all the loci tested (Supplementary Table 1), eight core regulators of endosomal trafficking had a conspicuous effect on mSpitz∷GFP localization (Fig. 1A-I). The functions of these loci are spread across the endosomal system with Rab5 and Vps34 controlling endosomal entry and maturation, while Vps23/TSG101, Vps25, Snf7A/Vps32 and Vps20 participate in sequential stages of multivesicular body formation, Carnation/Vps33 regulates Golgi to lysosome transport and Rab11 controls recycling and *de novo* movement to the apical surface. All exhibited either no or very few mSpitz∷GFP positive capes and absent or abnormal accumulation of mSpitz∷GFP in other compartments, whereas all exhibited wild-type Lamin C staining (i.e. abundant capes; not shown). These results indicate that a simple pathway for mSpitz∷GFP accumulation in capes that involves a diffusion and capture model following cotranslational import to the ER membrane is improbable. Instead it would appear that mSpitz∷GFP is first trafficked to the endosomal system (most likely *via* the plasma membrane), before arriving at the nuclear envelope. Evidence exists for trafficking of mSpitz to the apical plasma membrane followed by reinternalization and cleavage in endosomes, prior to basolateral secretion ^26^, and similar routing has documented for Wnt and Hedgehog ligands ^27^. However, the fact that uncleaved precursor travels to the perinuclear region *via* an endosomal route is a new observation, and may reflect a novel ‘salvage’ pathway to ensure that the immature form is not accidentally presented on the lateral membrane. Alternatively, there may be continuous circulation of mSpitz to facilitate a rapid signal generation by cleavage.

**Figure 1.**
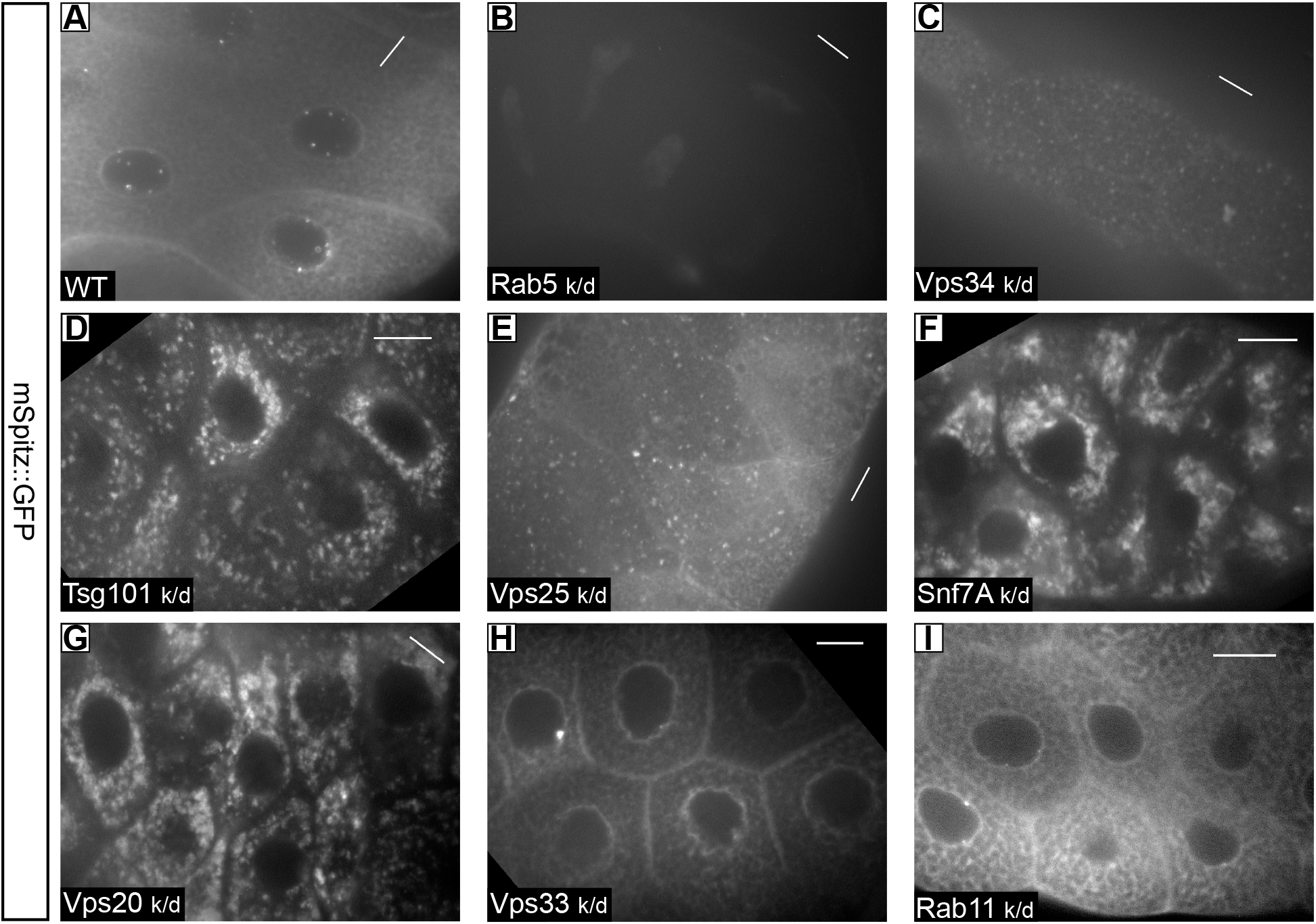
Endosomal regulators are required for mSpitz localization. Shown are low magnification views of salivary glands expressing mSpitz∷GFP along with UAS-RNAi constructs for the indicated genes: **A.** AB1-Gal4 > UAS-mSpitz∷GFP; **B.** AB1-Gal4 > UAS-Rab 5^RNAi^, UAS-mSpitz∷GFP; **C.** AB1-Gal4 > UAS-Vps34^RNAi^, UAS-mSpitz∷GFP; **D.** AB1-Gal4 > UAS-Vps23/TSG101^RNAi^, UAS-mSpitz∷GFP; **E.** AB1-Gal4 > UAS-Vps25^RNAi^, UAS-mSpitz∷GFP; **F.** AB1-Gal4 > UAS-Snf7/Vps32^RNAi^, UAS-mSpitz∷GFP; **G.** AB1-Gal4 > UAS-Vps20^RNAi^, UAS-mSpitz∷GFP; **H.** AB1-Gal4 > UAS-Carnation/Vps33^RNAi^, UAS-mSpitz∷GFP; **I.** AB1-Gal4 > UAS-Rab11^RNAi^, UAS-mSpitz∷GFP. Knockdown of these loci only, amongst all those tested (Supplementary Table 1), had an effect on mSpitz∷GFP distribution. However, none affected the appearance and frequency of cytoplasmic capes as indicated by Lamin C staining (not shown). Scale bars represent 20μm. The long axis of each scale bar indicates the direction of the proximal distal axis of the gland.

Interestingly, a retrograde pathway involving endosomes has also been identified in vertebrates for both the uncleaved and the membrane bound C-terminal fragment of Heparin-binding EGF-like growth factor ^7^, suggesting that this may be a conserved pathway. In this example the retrograde pathway appears to involve passage back through the Golgi and Rab6 mediated return to the ER ^6^. However, the close association of cytoplasmic capes with endosomes ^19^ suggests the possibility of direct transfer to the INM from the latter to the former, a pathway that was recently shown to exist ^8,9^. We previously showed that there is an enrichment of ubiquitylated proteins in cytoplasmic capes ^19^, and this strongly suggests that protein traffic from the endosomal system is being directly routed to this location prior to ubiquitin removal. Direct transfer from endosomes to the ONM would bypass the Golgi/ER.

Cytoplasmic capes in *Drosophila* salivary glands are not only associated with endosomes, but also with large granules and sites of perinuclear vesicle (PNV) formation ^19^. A recent report showed that PNV formation in the nuclei of muscle cells is part of the NCRE pathway for polyadenylated RNA’s and contain mega-RNP’s ^16,17^. A key regulator of this pathway is a cleaved C-terminal fragment of the Frizzled2 receptor (Fz2C), which enters the nucleus and is found at the export sites ^16^. In the supplementary data of Speese *et al*. (2012), this fragment was also shown to be present in salivary gland nuclei in structures that strongly resembled the cytoplasmic capes we previously described; however, RNA was not shown to be present at these locations in this tissue and so we sought to determine this.

Cytoplasmic capes can be readily identified by staining for Lamin C or by their accumulation of the immature EGF Receptor ligand mSpitz∷GFP ^19^. Fz2C puncta are distributed throughout the cytoplasm, but concentrate in the nucleoplasm and are prominent at sites of RNP export ^16^. Costaining 3^rd^ instar salivary glands for Fz2C and Lamin C showed strong colocalization (Fig. 2A, B), often at the base of the cape, where we previously showed that the PNV are most concentrated ^19^. These results confirm the observation of Speese *et al.* (2012) and strongly suggest that the cytoplasmic capes are associated with RNP export.

**Figure 2.**
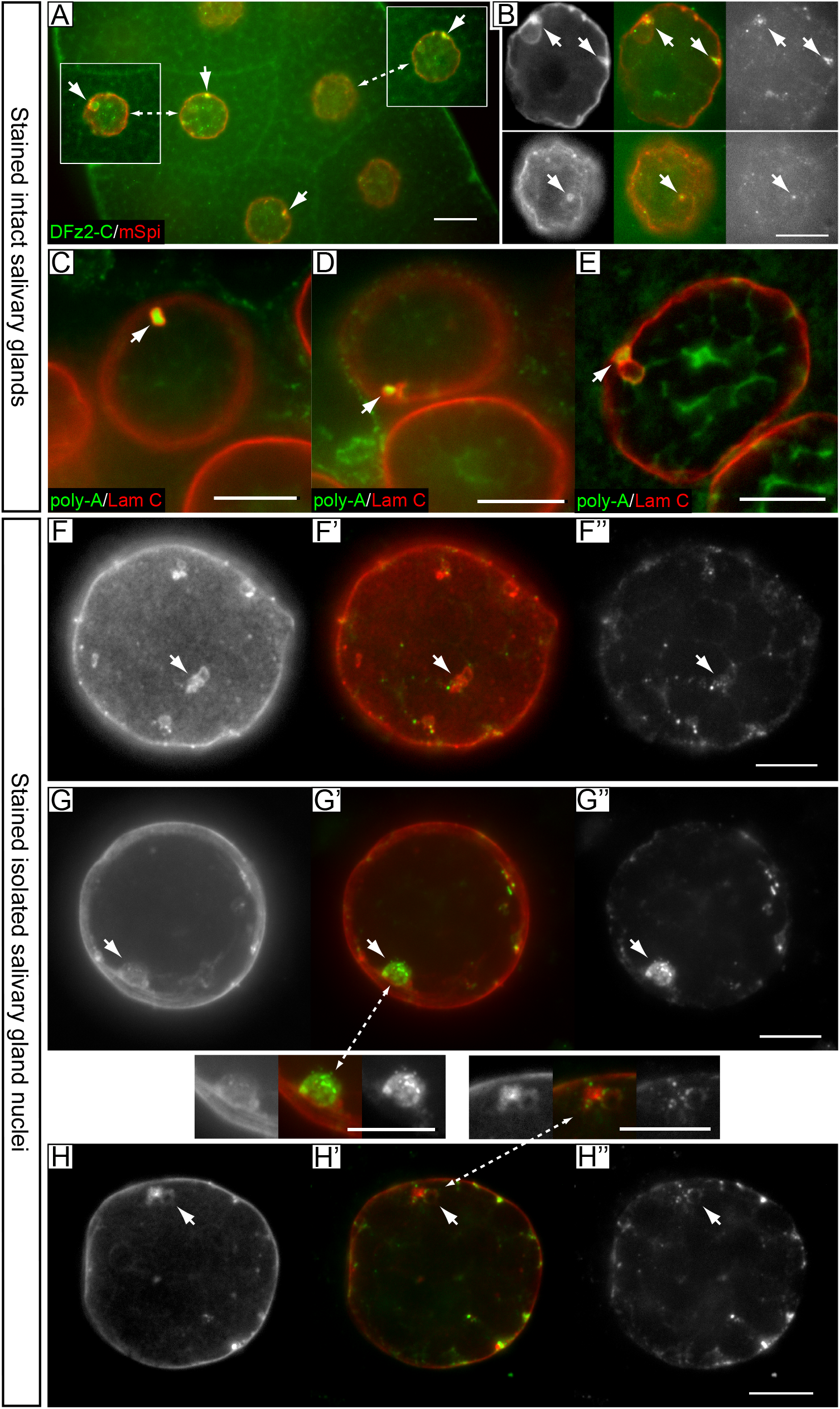
Cytoplasmic capes are strongly associated with RNP export sites. **A,B.** Nuclei in whole salivary glands stained for Fz2C (green) and Lamin C (red). A is a low magnification confocal section showing several nuclei. The insets show additional nearby planes of the indicated nuclei illustrating additional cape/Fz2C associations (arrows). B shows two focal planes in a single nucleus illustrating the conspicuous association of Fz2C puncta (RNPs) with capes (arrows). **C-E.** Three examples of *in situ* hybridization for polyA-containing mRNA (green) in intact salivary glands followed by costaining for Lamin C to reveal cytoplasmic capes. Colocalizations of mRNA in capes is indicated by the arrows. The success of these hybridizations is not high and mRNA signal is usually confined to the more superficial/exposed nuclei at the anterior of each gland. **F-H”.** Three examples of *in situ* hybridization for polyA-containing mRNA (green) in isolated salivary gland nuclei followed by costaining for Lamin C to reveal cytoplasmic capes. In each triptych Lamin C is shown to the left and RNA to the right of the merged central image. Examples of colocalization between RNA and capes is indicated by the arrows. Using this technique many RNP are detectable and almost all identifiable capes are associated with RNP. Multiple RNPs are often seen at the larger capes (enlargements between panels G and H). All images are single confocal planes except E, G and H, which are maximum projections of 3, 6 and 6 images (0.5μm interval), respectively to capture the whole cape. Scale bars represent 20μm in panel A and 10μm in panels B-H”.

We next looked directly for the presence of mRNA at the capes by *in situ* hybridization using an oligo-dT probe as previously described ^16^ followed by costaining for Lamin C. With low efficiency we identified mRNA-positive signals in some of the capes (Fig. 2C-E) but always in far fewer capes than we originally saw to be present in these nuclei ^19^, or that might be expected from our Fz2C costaining. It was conspicuous in our samples that *in situ* hybridization was most often seen where the nuclei were relatively superficial at the anterior of the gland, suggesting that probe penetration was a limiting factor in this tissue. We therefore modified the original protocol to gently disrupt the glands during fixation to release nuclei, which were recovered on slides by centrifugation for probing (see Materials and Methods). *In situ* hybridization could then proceed without the hindrance of surrounding cytoplasmic contents. The results of this protocol change are dramatic. Many RNP are detectable in each nucleus throughout the nucleoplasm, and we can now see that nearly every identifiable cape has a positive signal (Fig. 2F-H). In the best examples we are able to see multiple, individual polyA+ mRNA containing puncta in the large capes (insets between Fig. 2G,H). We conclude that cytoplasmic capes are very strongly associated with mRNA export. Capes vary dramatically in size ^19^ and the smallest (which are the majority) would likely not be resolved by light microscopy. In our previous ultrastructural characterization of the capes (*ibid*), even the small ones were associated with PNV, so we suspect that all capes are NCRE associated; however, it remains to be seen if all sites of NCRE are cape associated. Together, these data suggest that capes are sites of both import (endosome→nucleus) and export (RNP→cytoplasm) activities associated with the nuclear envelope. With so little currently known about either of these processes, we chose to further investigate the properties of cytoplasmic capes *per se*, and will pursue mechanistic studies on their relationship with the NCRE and endosomal transport in future publications.

Since membrane will also be transferred during both of these activities the local expansion of the nuclear envelope that creates a cape may well be linked to such transport. However, these are large structures and such a mechanism would only result in cape formation following many transfer events. This in turn predicts that capes should be persistent, perhaps growing structures. The presence of multiple RNP associated with cytoplasmic capes indicates that multiple rounds of PNV budding are indeed occurring at these sites (Figure 2G,H; ^19^). To begin to investigate this question we imaged capes in intact wandering 3^rd^ instar larvae at widely spaced time points to see if they are persistent and growing. To do this larvae were immobilized between dry agarose pads and a cover slip as previously described ^28^. Using the mSpitz∷GFP marker, capes were easily seen through the larval cuticle and could be followed for a period of about 90 minutes prior to pupation. As shown in the examples in Figure 3, capes persist for at least this period of time with little overt change in their size or morphology as judged by 3D reconstruction (see also supplementary movies S1-2). Speese *et al.* (2012) tracked RNP’s exiting the vicinity of the nucleus *via* the Fz2C-dependent pathway in just a few minutes. If this rate is typical of RNP export at all such sites this suggests that cape-associated RNP export events must persist through multiple rounds of budding and fusion at individual capes rather than assembling a cape *de novo* for each round of export. Similarly, the stable size of at least the larger capes is contrary to the hypothesis that on going transport leads to cape growth. However, we have only been able to do this imaging in wandering 3^rd^ instar larvae and it may be that transport is less active at this stage because the larvae are no longer growing (see below), so it remains possible that the initial growth of a cape is tied to membrane movement from the INM to the ONM and/or direct endosomal delivery of membrane.

**Figure 3.**
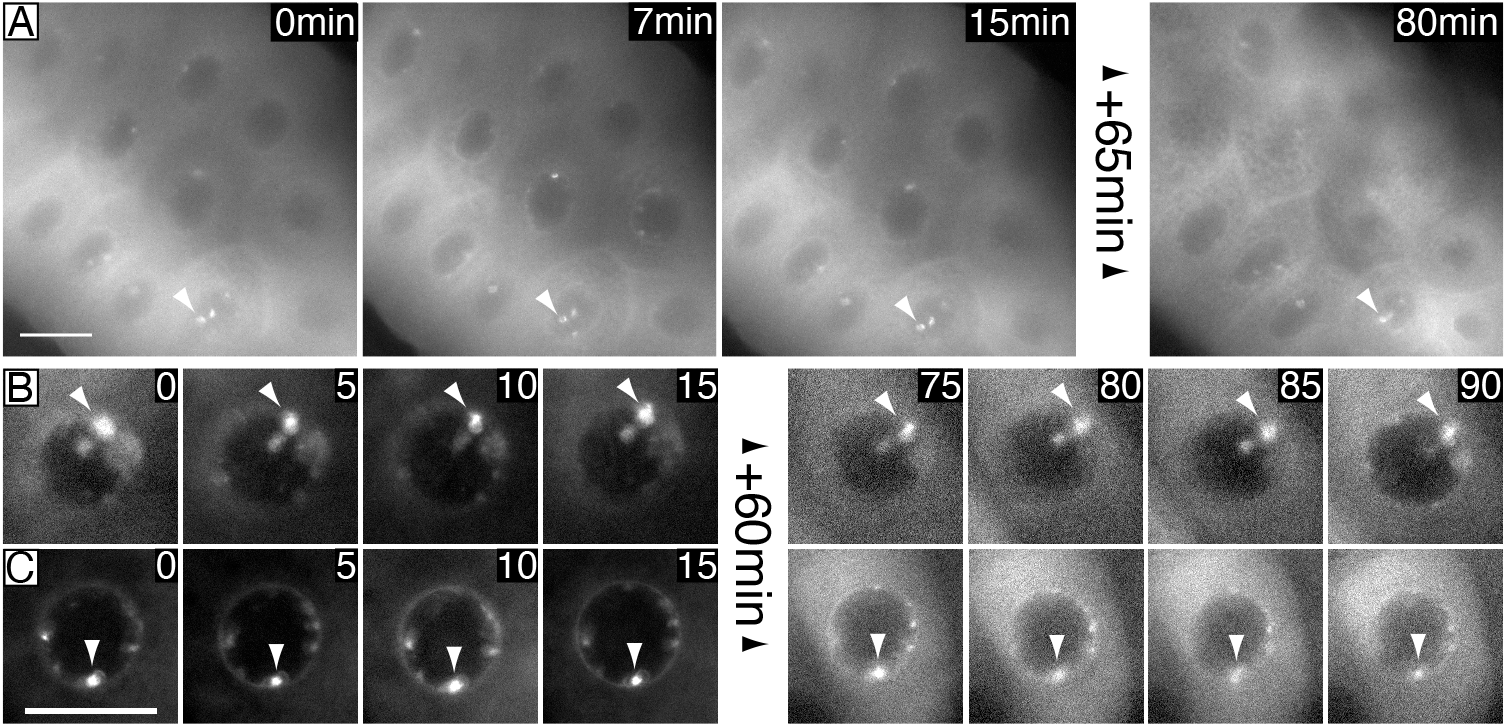
Cytoplasmic capes persist without growth for at least 90 minutes. All images show mSpitz∷GFP fluorescence imaged directly through the cuticle of living 3^rd^ instar larvae. Confocal image stacks were taken once a minute for 15 min, following which the larvae were released back to their medium. After one hour each larva was reimaged as before. **A.** Low magnification views of a gland showing the presence of many capes. Muscle activity in the larva distorts the gland unpredictably and so not all capes in the first frame are seen in subsequent frames. The presented focal planes were selected to follow the indicated capes (arrowhead). Cape persistence is the same in the other nuclei, which have moved out of focus. The initial 15 min sequence is presented as supplementary movie S1. **B-C.** Detail of capes in two nuclei followed for a total of 90 min. 90 minutes represented the maximum time we could image this stage because pupation and apoptosis follows shortly after. The signal to noise ratio was universally higher in the second sequence as the cells shrink due to glue secretion. The full sequence from C is presented in Supplementary movie S2. Scale bars represent 20μm.

Little is known about the developmental origin and timing of cytoplasmic capes. In their original description, capes were reported to be prevalent in the salivary glands of a number of *Drosophila* species during the 3^rd^ larval instar but the timing of their developmental appearance was not tightly defined ^20^. We therefore asked whether capes appeared at a specific time during larval development. To do this we utilized a Lamin∷GFP gene trap ^29^ as a marker and scored glands obtained from larvae of various ages raised at 25°C on standard media. Typically low-density cultures are used for such experiments to prevent density-dependent growth effects, ensuring that the size of the larvae closely parallels its developmental stage. We serendipitously discovered that this practice has the potential to obscure simple growth effects. Glands from moderate density cultures were dissected and immediately imaged without fixation. The results have been divided into four groups: Those with no visible capes (‘none’ in Fig. 4F), those with much less than 50% of nuclei exhibiting at least one cape (‘few’ in Fig. 4F), those with many nuclei (>50%) exhibiting capes (‘many’ in Fig. 4F) and a category we call ‘transitioning’ (‘trans’ in Fig. 4F). Transitioning larvae exhibited two conspicuously different Lamin∷GFP patterns within the same gland – smaller nuclei with a very uniform signal and slightly larger nuclei with more distinct outlines that often exhibited a few capes (Fig. 4A). In addition, we measured the diameter of each gland in a saggital focal plane midway along the proximal distal axis. Finally, a sampling of larvae were staged (1^st^, 2^nd^ or 3^rd^ instar) by examination of their mouthpart morphology (Fig. 4A’-E’; ^30^). Figure 4F shows the gland diameters within each of our four categories, rank ordered according to the mean diameter in each cape number category. To our surprise the presence or absence of capes does not seem to be determined by developmental stage but by gland size. The 2^nd^ instar larvae we imaged had no capes; however, it is clear that 3^rd^ instar larvae may have glands in any of the cape number categories.

**Figure 4.**
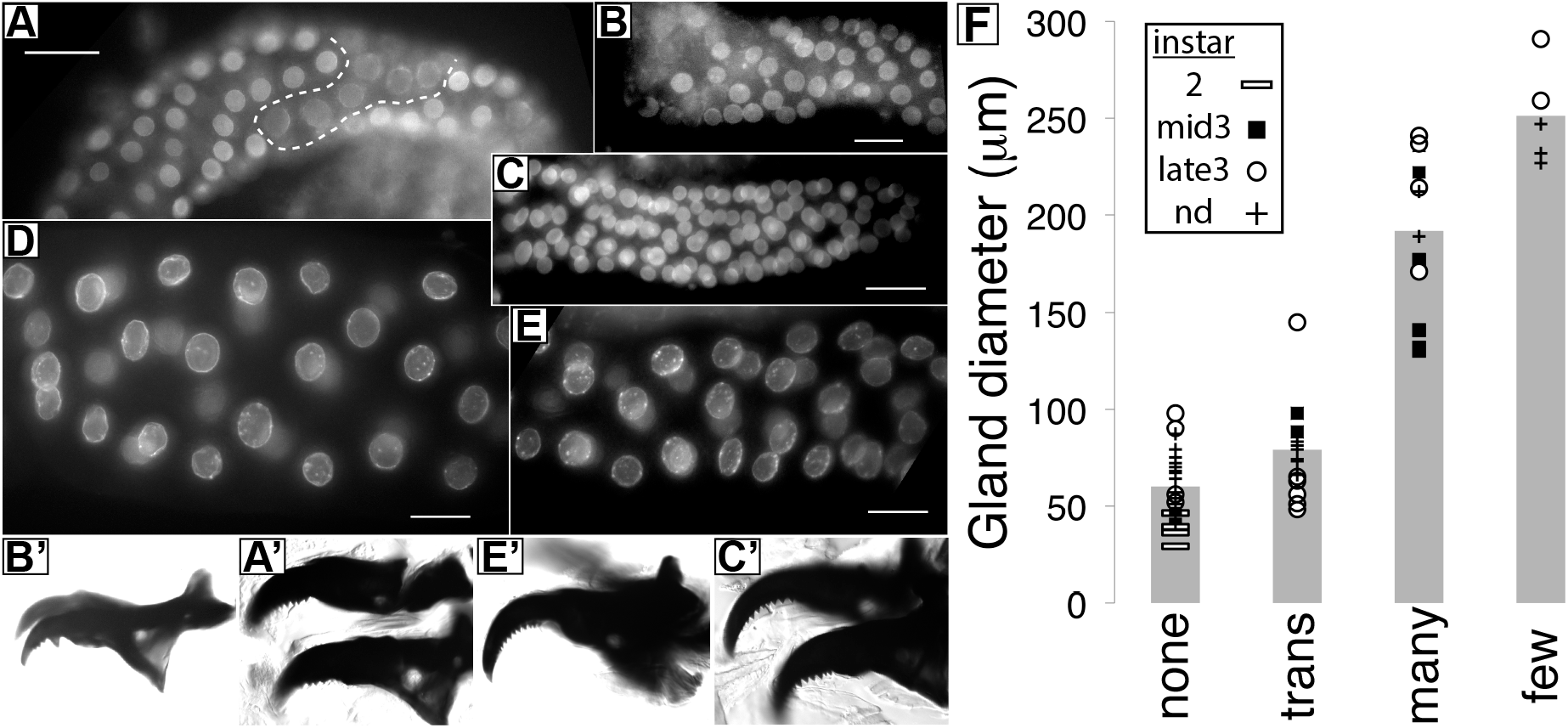
Cape appearance is a function of salivary gland size not age. **A-E** show salivary gland nuclei labeled by a Lamin∷GFP gene trap. Representative wide-field images are shown of glands from the ‘no nuclei with capes’ (B, C; ‘none’ in F), ‘transitioning’ (A, ‘trans’ in F) and >50% of nuclei with capes (D, E, ‘lots’ in F) categories. A’-C’ and E’ show corresponding mouthpart images used to stage each larvae (B’ is 2^nd^ instar; A’, C’ and E’ are all 3^rd^ instar). **F** Bar chart plotting the mean diameter of 81 glands from 44 larvae according to the categories of no nuclei with capes (‘none’), transitioning (‘trans’), >50% of nuclei with capes (‘many’), <<50% with of nuclei with capes (‘few’). Each gland had its diameter measured in a saggital plane and the categories are rank-ordered according to the mean for that category. Plotted points indicate the raw data. Where determined the developmental stage is indicated by the appropriate symbol: horizontal bars are second instar (3 teeth on mouthparts); circles indicate late 3^rd^ instar (9 teeth on mouthparts); and filled squares are 3^rd^ instar with intermediate tooth numbers (5-8 teeth on mouthparts). Plus signs indicate glands from larvae that were not staged. Scale bars represent 50μm.

The 3^rd^ instar is a time of massive tissue growth when larval mass increases several fold in just 3 days ^30^. Most of this mass is accrued without cell division, and the cells of most tissues expand their nuclear genome *via* endoreduplication to support the biosynthetic needs of this growth ^31,32^. Salivary gland cells develop to a huge size, a volume that is supported by a huge degree of endoreduplication, which may exceed 2000 × C ^33^, and is largely dedicated to the synthesis of glue proteins that will secure the pupa to it’s substrate. The ‘no capes’ category has a mean gland diameter of 60μm, while the ‘transitioning’ glands where some of the nuclei are slightly larger have a mean gland diameter of 79μm, and the glands with ‘lots’ of capes have a mean diameter of 192μm along with the classic giant nuclei of these organs. This suggests that capes appear concomitantly with the increasing biosynthetic activity of these cells, and we speculate that this is in response to an increased need for mRNP export. Interestingly, the ‘few’ category has larger glands (mean of 250μm), suggesting that as the glands reach their maximum size, just before mass secretion of the glue protein granules and apoptosis, that the number of capes may decline in parallel with a cessation of gene expression and glue protein export. It will be interesting in future studies to see if cape appearance closely parallels the onset of endoreduplication.

We previously showed that knockdown of β_H_-spectrin caused a major reduction in mSpitz∷GFP accumulation in capes ^19^. Comparison of Lamin C staining in wild-type and β_H_^RNAi^ glands reveals that knockdown of β_H_ drastically reduces the number and size of capes in salivary gland nuclei (Fig. 5A,B,D-E’), suggesting that our previous result reflected the absence of capes as a destination for mSpitz∷GFP. We also examined cape formation in larvae dependent upon a tetramerization deficient α-spectrin, *α*-*spec*^*R22S 34*^. *α*-*spec*^*R22S*^ caused an equally dramatic reduction in cape number (Fig. 5D, D’, F, F’) indicating that spectrin tetramerization is a requirement for cape formation. Tetramerization of α-spectrins and β-spectrins creates the F-actin crosslinking form of the protein, so this result also suggests that the F-actin cytoskeleton may be involved in cape formation.

**Figure 5.**
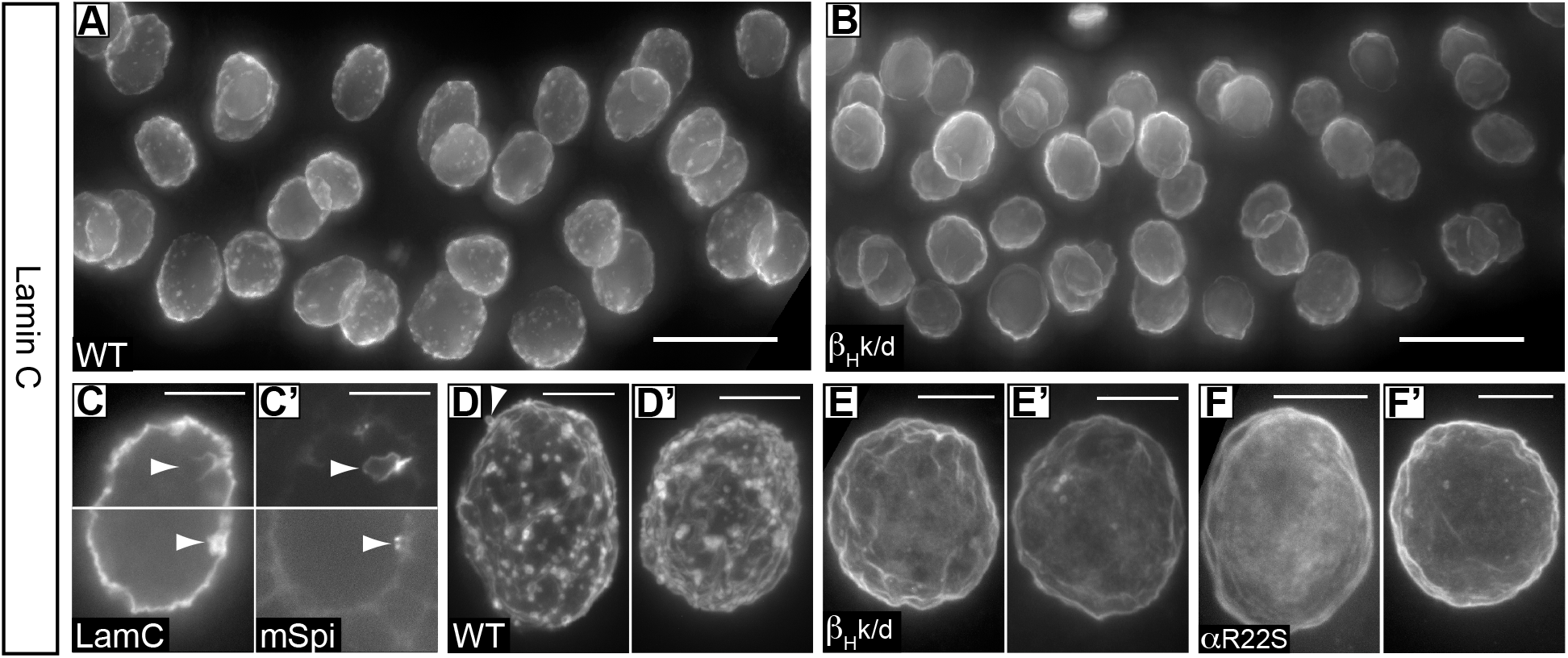
Cape formation is dependent on α- and β_H_-spectrin. Shown are low magnification views of gland nuclei stained for Lamin C or Lamin C and mSpitz∷GFP ( C,C’). **A,B**. Maximum projections of image stacks through AB1-Gal4 (driver only; WT) and AB1-Gal4 > β_H_^RNAi^ (β_H_k/d) salivary glands. Each wild-type nucleus has many capes, whereas each β_H_^RNAi^ nucleus has few if any capes. **C,C’** A nucleus costained for Lamin C (C) and mSpitz∷GFP (C’). Two different single image planes (top/bottom) are shown, each with a cape of different size (arrowheads). **D-F’** Two examples each of nuclei stained for Lamin C from AB1-Gal4 (driver only; WT), AB1-Gal4 > β_H_^RNAi^ (β_H_k/d) and *α*-*spec*^*R22S*^ (R22S) nuclei. Each is a maximum projection through a whole nucleus. Loss of β_H_ or disruption of spectrin tetramerization leaves very few capes, indicating that the absence of mSpitz∷GFP in this genotype ^19^ is likely due to the absence of capes. The arrowhead in D indicates an example of an outwardly oriented bleb/cape on the nuclear envelope. Scale bars represent 50μm in C and D and 10μm in D-F’.

β_H_ function has been closely associated with endosomal compartments ^35^, providing one possible route by which it could modulate cytoplasmic cape formation. However, spectrins have also been found inside the nucleus of vertebrate cells. βIVΣ5-spectrin is a predominantly nuclear isoform that associates with PML bodies ^36^ and the nuclear matrix ^37^. αIIΣ*spectrin is also found in association with the nuclear matrix (*ibid*) and has well-documented roles in DNA inter-strand crosslink repair mediated by the FANCA/C proteins in association with the XPF/XPG/XP complex ^38,39^. β_H_ has not yet been localized to the nuclear envelope or nucleoplasm by immunofluorescence; however, β_H_ along with the other spectrins were identified as a component of the nuclear matrix ^40^ suggesting that the role of β_H_ in cape formation and nuclear envelope biology could even be direct. Our identification of α-spectrin and β_H_ as key components for cytoplasmic cape formation suggests that there are additional roles for spectrin in nuclear biology that should be a fruitful subject for further study.

Interestingly, the original definition for the term ‘cytoplasmic cape’ was specifically applied to characteristic invaginations of both the INM and the ONM ^20^. However, it is clear from the early literature that first saw these structures that capes are one of a number of related structures all containing ‘oval bodies’ (i.e. RNP) where the INM and ONM may both bleb outwards from the nuclear envelope, or occasionally go in different directions ^41^ and the relative prevalence of these forms may depend on the species studied ^21,25,41–44^. While our previous serial block face imaging showed that the double invagination of the capes have been predominant in our samples ^19^, we do see outward blebs in some cases (arrowhead in Fig 5D). It is therefore a unifying observation that all of these morphologies disappear in the spectrin mutant backgrounds leaving a smooth nuclear envelope.

## Conclusions and perspective

We had previously shown that nuclear membrane infoldings, known as cytoplasmic capes, are closely associated with endosomal compartments ^19^. Here we show that capes do not depend on endosomes for their formation, but that at least one protein present in cape membranes (i.e. mSpitz) depends upon endosomal activities to accumulate at this location, thus emphasizing the close link between these membrane compartments. This report also documents a close association of non-canonical RNP export with cytoplasmic capes. The *Drosophila* salivary gland offers a unique opportunity to investigate these structures since they appear to become exceedingly large in these giant nuclei (see ^19^ for a comparison to wing disc capes), perhaps due to the intense biosynthetic activity in these giant cells. In line with this notion, our data suggests that these cape structures may be epicenters for multiple rounds of non-canonical RNP export as well as protein delivery from endosomes. It is currently unknown whether cape formation is a cause, consequence or an independent and parallel process to endosomal delivery and/or RNP export. Also, the role of mSpitz accumulation in this specific nuclear subcompartment and its relationship to EGF Receptor signaling is currently unknown. Future investigations will probe these possibilities.

## Materials and Methods

### Fly Strains

Oregon-R or yellow white stocks were used as wild-type strains. The AB1-Gal4 driver (y^1^ w*; P{w^+mW.hs^=GawB} AB1) was obtained from the Bloomington Stock Center (Indiana University, Bloomington, IN; BDSC Cat# 1824, RRID:BDSC_1824). UAS-mSpitz∷GFP was gift from Dr. Eyal Schejter (Weitzman Institute, Rehovot, Israel). UAS-β_H_^RNAi^ and the generation of *α*-*spec*^*R22S*^ rescued flies were as previously described ^34,35^. UAS-RNAi lines are listed in Supplementary Table 1 and were acquired from the Vienna *Drosophila* Resource Center (VDRC; Campus Vienna Biocenter, A-1030 Vienna, Austria; RRID:SCR_013805). For homozygous viable UAS-RNAi inserts the genotypes imaged were either UAS-RNAi/+; AB1-Gal4/+ or UAS-RNAi/AB1-Gal4. The UAS-TSG101^RNAi^ line (VDRC #23944) is homozygous sterile and was therefore introduced from a stock with a marked balancer (CyO, Act-GFP) and only non-fluorescent larvae were dissected. The Lamin∷GFP genetrap line was obtained from Dr. Melissa Rolls (The Pennsylvania State University, University Park, PA).

### Fluorescent *In Situ* Hybridization (FISH)

#### FISH on intact salivary glands from wandering 3^rd^ instar larvae

Glands were dissected in cold PB+RVC (100 mM Na Phosphate Buffer pH 7.2, 10 mM Ribonucleoside-vanadyl complex; New England Biolabs, Ipswich, MA). During dissection, glands were kept in ice cold PB+RVC for no more than 30 min to minimize RNA degradation. Glands were then pre-extracted in PB+RVC with 0.5% TritonX-100 for 10 min on ice, then fixed in 4% Paraformaldehyde (PFA) in 100 mM Na Phosphate Buffer pH 7.2, for 30 min at 4°C, and immediately postfixed in cold 100% Methanol for 10 min at 4°C. Glands were then re-hydrated through a series of 10 min washes in 70%, 50%, 30%, 0%, and 0% Methanol in 0.2% PB+RVC with 0.2% TritonX-100 at room temperature (Rm T) and equilibrated in Hybridization Buffer (2X SSC, 10% dextran sulfate, 20 mM RVC, 15% formamide) by incubating in 50% Hybridization Buffer:PB+RVC plus 0.2% TritonX-100 for 5 min, then 100% Hybridization Buffer + 0.1 mg/ml yeast tRNA for 10 min at RmT. Hybridization Mixture was prepared as follows: An appropriate volume of Probe Mix (5.26ng/μl 5’Digoxigenin-polyT(24)-3’Digoxigenin [Integrated DNA Technologies, Inc., Coralville, IA], 52.6 ng/μl sheared salmon sperm DNA [Rockland Immunochemicals, Inc., Limerick, PA]) in 31.6% deionized formamide was heated to 80°C for 10 min, then immediately cooled on ice. 2X Hybridization Buffer (4X SSC, 20% dextran sulfate, 40 mM RVC), 20 mg/mL yeast tRNA, and 50X Denhardt’s Reagent were added to yield final concentrations of 2.5 ng/uL probe, 25 ng/uL sheared salmon sperm DNA, 15% formamide, 1X Hybridization Buffer, 0.1 mg/mL yeast tRNA, and 1X Denhardt’s Reagent. Glands were transferred to the Hybridization Mixture and incubated for at least 18 hours at 37°C with gentle agitation. Following hybridization, glands were washed twice in 2X SSCt (SSCt=1X or 2X or 4X SSC + 0.001% TritonX-100) with 15% Formamide, twice in 2X SSCt alone, and twice in 1X SSCt for 15 min each. Glands were then incubated in sheep anti-Digoxigenin IgG (1:200; Sigma-Aldrich Cat# 11093274910, RRID:AB_2734716) in 4X SSCt overnight at 4°C. Glands were then washed once in 4X SSCt, once in 4X SSC with 0.1% Triton X-100, and twice in 4X SSCt for 30 min each. Glands were again fixed with 4% PFA for 10 min at RT, and subsequently washed four times for 15 min at RmT in 0.2% PBT (100 mM Phosphate Buffer pH 7.2, 0.2% TritonX-100). Glands were then incubated in donkey anti-Sheep IgG:FITC (1:200; Jackson ImmunoResearch Labs Cat# 713-095-147, RRID:AB_2340719) in 0.2% PBT overnight at 4°C. Following incubation, glands were washed four times for 30 min at RmT in 0.2% PBT, and then incubated in rabbit anti-FITC IgG (1:700; Thermo Fisher Scientific Cat# ANZ0202, RRID:AB_2536401), 1:30 mouse anti-Lamin C (developed by P.A. Fisher at the MPI for Biophysical Chemistry, Goettingen FRG, was obtained from the Developmental Studies Hybridoma Bank, created by the NICHD of the NIH and maintained at The University of Iowa, Department of Biology, Iowa City, IA 52242; DSHB Cat# lc28.26, RRID:AB_528339) in 0.2% PBT overnight at 4°C. Following incubation, glands were washed four times 30 min at room temperature in 0.2% PBT, and then incubated in goat anti-Rabbit IgG:Alexa488 (1:250; Thermo Fisher Scientific Cat# A-11034, RRID:AB_2576217) and goat anti-Mouse IgG:Alexa594 (1:250; Thermo Fisher Scientific Cat# A-11032, RRID:AB_2534091) in 0.2% PBT overnight at 4°C. Glands were then washed four times 30 min in 0.2% PBT, and then equilibrated in Mounting Medium (80% glycerol, 0.1 mM Tris, pH 8.5) for at least 1 hour before mounting and imaging.

#### For FISH on isolated nuclei

Salivary glands were dissected from ~25 wandering 3rd instar larvae in cold PB+RVC and held on ice for no more than 30 min to minimize RNA degradation. Glands were transferred in 250μl of PB+RVC to an ice cold 2mL Dounce homogenizer, combined with 250μL of 8% PFA in PBS (10 mM NaPO_4_, 130 mM NaCl pH7.2), and immediately homogenized by 10 strokes of a tight pestle. Homogenized glands were allowed to fix on ice for a total of 10 min, then 500μL of 1M Tris pH 7.5, 10mM RVC was added directly to the homogenizer. The homogenate was centrifuged through a 1mL sucrose cushion (10.3% (w/v) Sucrose, 100 mM Na Phosphate Buffer pH 7.2, 10 mM RVC) at 4°C at 1500xg in a swinging bucket rotor to remove most cellular debris. The pellet was resuspended in PB+RVC, spun onto poly-D-lysine coated microscope slides using a Cytospin centrifuge (Thermo Scientific, Waltham MA), and fixed to the slides by spinning 4% PFA though the Cytospin chambers. Slides were removed from the Cytospin and nuclei were fixed with methanol for 10 min on ice. All subsequent incubations were done on the slide where rehydration, hybridization, washing and imaging was the same as for the whole glands.

### Salivary gland dissection

To image nuclei in freshly dissected salivary glands, larvae were removed from food, rinsed, dissected in PBS, and imaged in less than 10 min. For staging, larval mouthparts were saved in ethanol and the surrounding tissue cleared in methyl salicylate prior to imaging.

### Larval Immobilization for timelapse imaging

*Drosophila* larvae were mounted on a microscope slide using a dried agarose pad, and a coverslip was taped in place to secure the larvae as previously described ^45^. Imaging was conducted immediately for 15 minutes. Larvae were released from the agarose pad by applying a drop of water before transfer to standard media between imaging sessions.

### Antibodies and Immunofluorescence

Primary antibodies used for immunofluorescence in this study were mouse anti-Lamin C (1:30; developed by P.A. Fisher at the MPI for Biophysical Chemistry, Goettingen FRG, was obtained from the Developmental Studies Hybridoma Bank, created by the NICHD of the NIH and maintained at The University of Iowa, Department of Biology, Iowa City, IA 52242; DSHB Cat# lc28.26, RRID:AB_528339); rabbit anti-DFz2-C (1:500; ^46^; was a gift from Dr. Vivian Budnik, University of Massachusetts Medical School, Worcester, MA 01605). Secondary antibodies for immunofluorescence in this study were Goat anti-Rabbit IgG Alexa Fluor 488 (1:250; Thermo Fisher Scientific Cat# A-11034, RRID:AB_2576217) and Goat anti-Mouse IgG Alexa Fluor 594 (1:250; Thermo Fisher Scientific Cat# A-11032, RRID:AB_2534091)

Immunofluorescence of larval salivary glands followed our previously published methodology ^35^. In brief, wandering 3^rd^ instar larvae were collected, dissected in PBS, stored in PBS on ice for no more than 10 min, and fixed for 20 min in 4% PFA on ice. Glands were washed in PBS [4X 15 min] and blocked for 1 hour in blocking solution (10% NGS, 0.5% Triton X-100, 0.3% deoxycholate, 0.2% saponin, 1X PBS) at room temperature. All subsequent washes and incubations were in Incubation Solution (5% NGS, 0.5% Triton X-100, 0.3% deoxycholate, 0.2% saponin, 1X PBS). Glands were incubated in 1° antibody (at each specified concentration) overnight at 4°C, washed 4 × 30 min, incubated in 2° antibody (1:250) overnight at 4°C, washed as before, then equilibrated in mounting medium prior to mounting and imaging.

### Imaging and image processing

Samples were imaged on a CARV II spinning disc confocal (BD Biosystems, Rockville, MD) with a Retiga EXi camera (Q Imaging systems, Surrey, BC) and iVision 4.5 software (Biovision, Exton PA). Post acquisition processing was limited to contrast stretching, except for Figure 1E and 2B where rolling ball background subtraction (45 and 300 pixel diameter respectively) was applied to decrease background haze and uneven illumination. To create the supplementary movies manually selected confocal planes at each timepoint were combined in a stack, aligned in Fiji using conspicuous fiduciary points with the TrackEM2 function. The aligned TIFF files were converted to movie sequences using iVision. Final processing and figure assembly for this paper was done using Adobe CS4 (Adobe, San Jose, CA).

## Acknowledgments

The authors would like to acknowledge James Ashley for helpful advice when getting started with the *in situ* hybridization procedure, Megan Radyk for supplying the *α*-*spec*^*R22S*^ larvae, Vivian Budnik for gifting the DFz2-C antibody, Eyal Schejter and Melissa Rolls for fly stocks, and Daniela Zarnescu for critically reading an early draft of the manuscript. This work was funded by NSF grant #112013 to CMT.

## Supporting information captions

**Supplemental Movie 1. Timelapse videos of cytoplasmic cape behavior in salivary gland cells of intact larvae.**

Live AB1-Gal4>mSpitz∷GFP third instar larvae expressing were restrained as described in the methods section, and imaged through the cuticle. This sequence covers 15 minutes. Cape size, intensity and number does not detectably vary. Stills from this video were used in Figure 3A.

**Supplemental Movie 2. Timelapse videos of cytoplasmic cape behavior in salivary gland cells of intact larvae.**

Live AB1-Gal4>mSpitz∷GFP third instar larvae expressing were restrained as described in the methods section, and imaged through the cuticle. In this video one nucleus was abstracted from a higher magnification sequence, aligned in ImageJ to reduce displacements due to nearby muscle activity. Two sequences are abutted each representing 15minutes. There was a 60minute gap between the two periods during which the larva was released back into food. The transition to the second sequence is conspicuous from the dramatic increase in background fluorescence in the cytoplasm, which we have seen in all such experiments and suspect originates late developmental changes in the gland just prior to pupation. Individual frames from this sequence were used in Figure 3C.

**Supplementary Table 1.** List of endosomal proteins targeted by RNAi and their effects.

List of endosomal proteins targeted by RNAi and their effects
| Compartment/<br>complex | Protein knocked<br>down | Effect on mSpitz<sup>1</sup><br>(mSpitz::GFP) | VDRC stock<br>number |
| --- | --- | --- | --- |
| **Apical delivery and slow recycling** |  |  |  |
| Recycling<br>endosome | Rab 11 | + | 108382 |
| **Early endosome entry and maturation** |  |  |  |
| Early endosome | Rab5 | +++ | 103945 |
|  | Vps34 | ++ | 100269 |
| **Fast recycling** |  |  |  |
| Early endosome | Rab4 | - | 106651 |
| **Late endosome / Multi vesicular body formation** |  |  |  |
| ESCRT I | Vps23/TSG101 | +++ | 23944 |
|  | Vps37A/Modr<sup>2</sup> | - | 104530 |
|  | Vps28 | - | 1894 |
| ESCRT II | Vps25 | ++ | 108105 |
|  | Vps22 | - | 110350 |
|  | Vps36 | - | 107417 |
| ESCRT III | Snf7A/Vps32 | +++ | 106823 |
|  | Vps20 | +++<sup>3</sup> | 26387 |
|  | Vps24 | - | 100295 |
| **Maturation to / fusion with lysosome** |  |  |  |
| Late endosome/<br>Lysosome | Rab7 | - | 40338 |
| HOPS/CORVETE | Car/Vps33 | + | 4548 |
|  | Dor/Vps18 | - | 107053 |

